# Wireless Neuromodulation at Submillimeter Precision via a Microwave Split-Ring Resonator

**DOI:** 10.1101/2022.07.22.501150

**Authors:** Carolyn Marar, Ying Jiang, Yueming Li, Lu Lan, Nan Zheng, Chen Yang, Ji-Xin Cheng

**Affiliations:** Department of Biomedical Engineering, Boston University, Boston, MA, USA; Graduate Program for Neuroscience, Boston University, Boston, MA, USA; Department of Mechanical Engineering, Boston University, Boston, MA, USA; Department of Electrical & Computer Engineering, Boston University, Boston, MA, USA; Division of Materials Science and Engineering, Boston University, Boston, MA, USA; Department of Chemistry, Boston University, Boston, MA, USA

**Author notes:** C. Marar and Y. Jiang contributed equally to this work. Department of Biological Engineering, Massachusetts Institute of Technology, Cambridge, MA, USA.

## Abstract

Microwaves, with wavelengths on the order of millimeters, have centimeter-scale penetration depth and have been shown to reversibly inhibit neuronal activity. Yet, microwaves alone do not provide sufficient spatial precision to modulate target neurons without affecting surrounding tissues. Here, we report an implantable split-ring resonator (SRR) that generates a localized and enhanced microwave field at the gap site with submillimeter spatial precision. The SRR breaks the microwave diffraction limit and greatly enhances the efficiency of microwave inhibition. With the SRR, microwaves at dosages below the safe exposure limit are shown to inhibit neurons within 1 mm from the gap site. Application of the microwave SRR to suppress seizures in an *in vivo* model of epilepsy is demonstrated.

## Introduction

Neuromodulation is a rapidly expanding field which has applications in neuroscience research, disease diagnosis, and treatment. Implantable neuromodulation devices are seeing greater use in the clinic for the treatment of conditions such as depression, epilepsy, and chronic pain [1, 2, 3]. Of these techniques, deep brain stimulation (DBS) is the most widely used, delivering electrical current via an implanted electrode to deep brain regions. The electrode, however, must be tethered to a subcutaneously implanted stimulator [4, 5, 6]. This requirement makes the device highly invasive, as surgery is required to change the implanted stimulator battery.

Electromagnetic waves, such as radio-frequency waves, have been used to non-invasively modulate various biological systems [7, 8, 9]. For example, transcranial direct current stimulation (tDCS) [7] and transcranial magnetic stimulation (TMS) [9] have successfully reached the deep brain to treat Parkinson’s Disease, depression, and epilepsy. However, due to the long wavelength (tens of meters) of the electromagnetic waves employed, tDCS and TMS offer poor spatial resolution of a few centimeters. Photons have sub-micron wavelength and provide single-cell modulation through optogenetics [10, 11]. Yet, the strong tissue scattering prevents photons from noninvasively reaching deep tissue. More recently, optical fiber-based optoacoustic neural stimulation has demonstrated sub-millimeter spatial resolution [12], but the need for coupling the laser to the fiber based optoacoustic emitter prevents wireless implementation.

Microwaves, with frequencies between 300 MHz and 300 GHz, fill the gap between optical waves and magnetic waves, yet have rarely been explored for neuromodulation. Microwaves have much longer wavelengths than photons and have been known to provide >50 mm penetration depth into the human brain noninvasively, while maintaining more than 50% of their energy [13, 14]. Microwave wavelengths are also much shorter than those of magnetic waves, promising higher spatial resolution for regional targeting. Reports of using the non-thermal effect of microwaves to modulate neural activity date back to the 1970s, where low intensity microwave was applied to Aplysia pacemaker neurons for extended time periods (> 60 s), and a reversible reduction in the firing rate was observed [15]. The mechanism was attributed to microwave perturbation of current flow inside axons. Since then, several studies have focused on the effect of chronic exposure to microwaves from cell phones, Wi-Fi, and other communication apparatus [13, 16, 17, 18, 19]. However, these studies utilized broadcasted microwaves, which lack spatial precision. Furthermore, the extended exposure time increases the risk of thermal damage to both targeted and surrounding tissues.

Here, we report minimally invasive microwave neuromodulation at an unprecedented spatial resolution by taking advantage of an implantable split-ring resonator (SRR) design. The SRR has a perimeter of approximately one half of the resonant microwave wavelength, thus acting as a resonant antenna. It couples the microwave wirelessly and concentrates it at the gap, producing a localized electrical field. We report the localization of a microwave field to >1 mm in space around the gap of the SRR via resonance with the microwave SRR at 2.1 GHz frequency. The device allows for neuromodulation with a submillimeter spatial resolution, beyond the microwave diffraction limit, while using a total dosage of ~500 J/kg, over 7 times lower than the threshold for safe microwave exposure [20]. We demonstrate the capability of the microwave SRR to inhibit neuronal activity transcranially and with submillimeter spatial precision. Additionally, we explore a potential application of the microwave SRR in an *in vivo* model of epilepsy.

## Results

### The SRR efficiently concentrates microwaves at the gap site

The SRR can be modeled as an LC resonance circuit where the ring acts as an inductor *L* and the gap acts as a capacitor *C*. When the SRR resonates with the incident microwave, a strong electric field is concentrated at the capacitor. To theoretically verify the resonance effect of the SRR and determine the resonant frequency, we applied finite element modeling of a copper SRR with a diameter of 2.54 mm in bulk water under electromagnetic fields from 0.1 GHz to 3 GHz. At 2.0 GHz, a strong electromagnetic field was observed at the gap site with intensity approximately 200 times larger than that of the surrounding medium (**Figure 1a**). The full width at half maximum (FWHM) of the microwave intensity at the gap was 0.45 mm (**Figure 1b**), whereas the wavelength of the microwave in water was ~17 mm.

**Figure 1:**
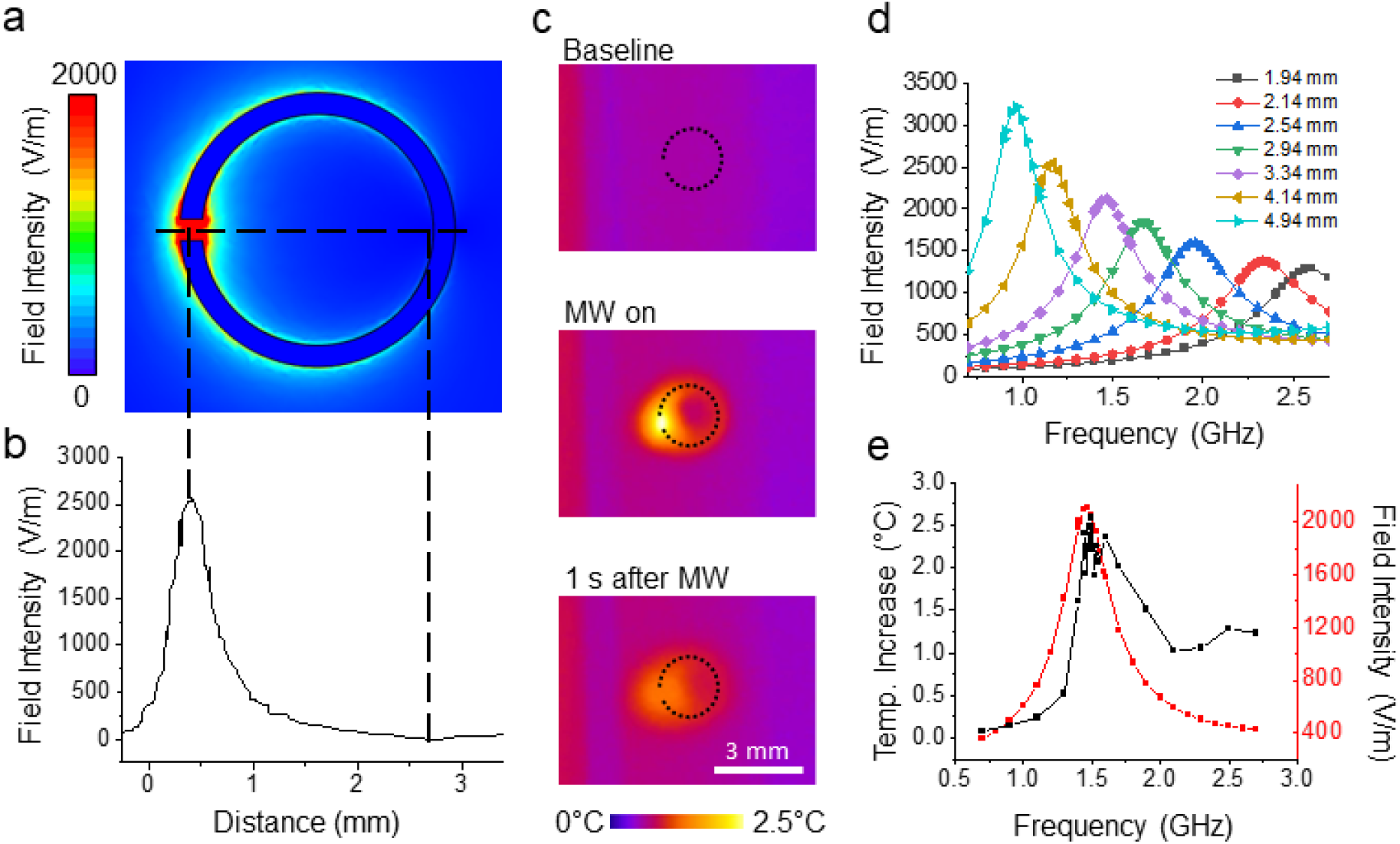
The SRR concentrates microwaves at its gap. (a) Simulated electric field intensity of a copper SRR with diameter 2.54 mm in bulk water under 2.0 GHz microwave irradiation and (b) profile of field intensity cross section; (c) thermal images of 2.54 mm diameter SRR before, during, and after 1 s microwave irradiation at 2.0 GHz and 2 W/cm^2^ demonstrating hotspot formation at the gap; (d) maximum temperature change at SRR gap and simulated microwave intensity for given frequencies for SRR with diameter 3.34 mm; (e) simulated electric field intensity at given frequencies for SRRs of varying diameter.

To experimentally validate hotspot formation, a copper SRR (outer diameter 2.54 mm, gap 0.2 mm, height 0.2 mm and width 0.2 mm) was fabricated through laser cutting. The SRR was covered in epoxy and placed at the air-water interface and imaged with a thermal camera (**Supplementary fig. 1a**). Microwave was delivered through a waveguide with the magnetic field perpendicular to the SRR plane at 2 W/cm^2^ and 2.0 GHz for 1 s. Thermal images shown in **Figure 1c** provide visual evidence of the hot spot at the SRR gap. A maximum temperature increase of 2.51°C was observed at the gap site. At off-resonance frequencies, the maximum temperature increase was less than 0.13°C. Such a hot spot at the resonant frequency confirms a localized, enhanced electric field predicted by our simulation.

To demonstrate the tunability of the resonant frequency, SRRs with varying diameters were simulated. The resonant frequency in an LC circuit is defined as 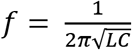. Increasing the perimeter of the SRR increases the inductance and consequently decreases the resonant frequency. In accordance, our simulation shows that by varying the diameter, we can tune the resonant frequency of the SRR (**Figure 1d**). To validate the simulation, an SRR with a diameter of 3.34 mm was irradiated with 2 W/cm^2^ microwave at frequencies ranging from 0.7 – 2.7 GHz. This SRR also achieved a maximum temperature increase of ~2.5°C with a peak at 1.5 GHz, in accordance with the resonant frequency determined by the simulation (**Figure 1e**). Collectively, these data suggest that the SRR generates an amplified electromagnetic field at the gap site with tunable resonant frequency.

For the neuromodulation applications presented here, we chose to use SRRs with resonant frequencies of 2.0 to 2.1 GHz, corresponding to wavelengths of 15 to 14.3 cm, respectively. These frequencies are at the low end of the dielectric loss spectrum for water, meaning they have low absorption in water. This is essential for achieving high penetration depth with minimal thermal damage in biological tissues. Larger wavelengths, although they have lower absorption, were not used in order to minimize the size of the implant.

### Microwave inhibits neuronal activity via a nonthermal mechanism

Microwave inhibition of neuronal firing through a non-thermal mechanism has been previously demonstrated in Aplysia pacemaker neurons [15] and avian neurons [19]. To verify that the inhibitory effect also occurs in mammalian neurons, we exposed cultured primary cortical mouse neurons to a microwave field at 1.0 GHz and 2 W/cm^2^ for 3 s (**Figure 2a)**. Neuronal activity was visualized by calcium imaging of GCaMP6f transfected neurons. Neurons exhibited spontaneous activity without microwave treatment. Immediately after microwave irradiation, neurons showed reduced firing rates for up to 50 s, after which they resumed their spontaneous firing patterns. This result confirmed direct microwave radiation at 2 W/cm^2^ can inhibit neuron activities and the inhibition is not induced by damage of the neurons. Neuronal activity was further quantified by area under the curve (AUC) of the normalized GCaMP6f fluorescence signal (F/F_0_). Almost all the neurons exhibited a decrease in activity and an average of 45.7% reduction was observed for 3 s of microwave irradiation (**Figure 2b**). Similar inhibition effects were observed at frequencies ranging from 0.7 – 2.4 GHz. Notably, the neurons with the highest baseline activity exhibited the greatest changes. Simultaneous thermal imaging of the cell culture under microwave irradiation shows that the temperature increase during 3 s microwave exposure was 1.6°C (**Figure 2c**), which is known to have no significant modulation effect in mammalian neurons [21, 22]. These results confirm the capabilities of microwaves to inhibit mammalian neurons via a non-thermal mechanism.

**Figure 2:**
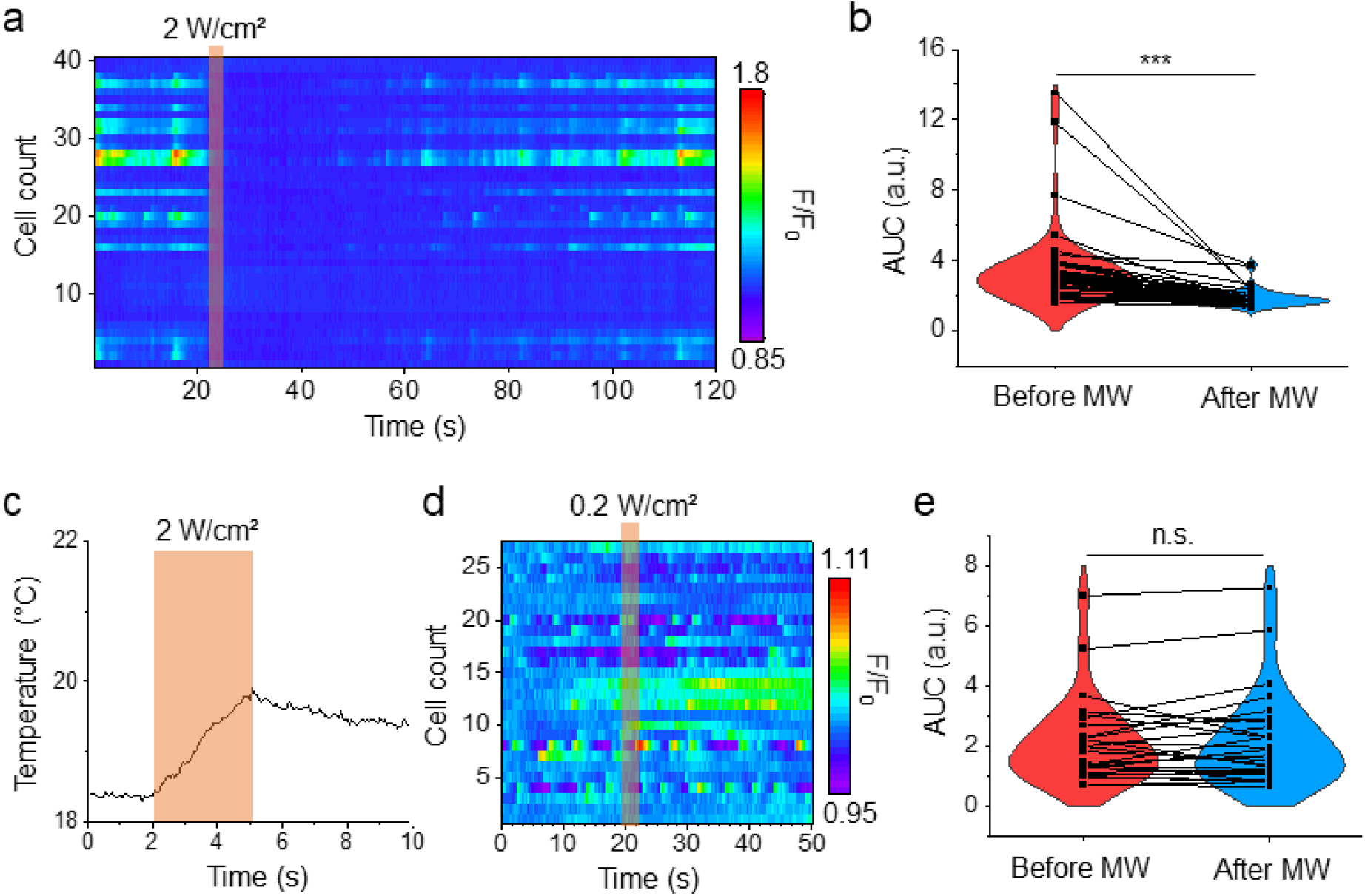
Microwaves inhibit neuronal activity. (a) GCaMP6f fluorescence heatmap for neurons under direct 2 W/cm^2^ microwave irradiation at 1.0 GHz for 3 s; (b) AUC of F/F_0_ traces for cells in (a) before and after microwave irradiation (n=49); (c) thermal change in cell culture medium at the SRR gap site during 3 s of 2 W/cm2 microwave irradiation at 1.0 GHz; (d) GCaMP6f fluorescence heatmaps for neurons under direct 0.2 W/cm^2^ microwave irradiation at 2.0 GHz for 3 s; (e) AUC of F/F_0_ traces for cells in (d) before and after microwave irradiation (n=27). Orange shaded boxes indicate microwave on; statistical significance calculated using paired sample Wilcoxon signed rank test where ***p<0.001, **p<0.01, *p<0.05, n.s. not significant.

To test the capabilities of microwaves to inhibit neurons at a lower power density, neurons were exposed to 0.2 W/cm^2^ microwave for 3 s at 2.0 GHz (**Figure 2d**). At this power density, no significant changes in neuronal activity were observed (**Figure 2e**). Therefore, microwave alone is not able to significantly reduce neuronal activity at a low power density. This means that in order to use microwave radiation for neuromodulation with improved safety, the efficiency of inhibition must be improved.

### Microwave SRR inhibits neurons with improved efficiency and submillimeter spatial precision

For safe application *in vivo*, it is desirable to minimize the required microwave dosage for inhibition. The confinement of microwaves to the SRR gap creates a concentrated electric field at the gap site. Thus, we expect the inhibition effect of microwaves to be enhanced in the presence of the SRR, reducing the required microwave dosage and limiting the affected area. The copper SRR with a 2.54 mm diameter and resonant frequency at 2.0 GHz was used for subsequent experiments. To investigate how the SRR could enhance the efficiency and spatial precision of microwave inhibition, the SRR was submerged in the culture medium above the primary cortical neurons at a distance of ~100 μm from the cells (**Supplementary fig. 1b**). The SRR was oriented perpendicular to the culture dish and microwave was delivered with H field perpendicular to the SRR plane.

To investigate the dependence of microwave inhibition on power density, the SRR was irradiated with 2.0 GHz microwave for 3 s at power densities of 2.0 W/cm^2^ (**Figure 3a**), 1.0 W/cm^2^ (**Figure 3b**), 0.2 W/cm^2^ (**Figure 3c**), and 0.02 W/cm^2^ (**Figure 3d**). Ordering neurons by distance from the SRR gap in heatmaps demonstrates the dependence of spatial resolution on power density. As power density decreases, cells further from the gap experience smaller changes in fluorescence intensity. To quantify the dependence of inhibition efficiency on power density, AUC of F/F_0_ was calculated for 15 s periods immediately before and immediately after microwave treatment. Activity in neurons within 1 mm from the SRR gap was reduced by averages of 51.8%, 36.2%, 36.3%, and 13.9%, for power densities of 2.0 W/cm^2^, 1.0 W/cm^2^, 0.2 W/cm^2^, and 0.02 W/cm^2^, respectively (**Figures 3e-h)**. These results indicate that successful inhibition was observed at 2.0 W/cm^2^, 1.0 W/cm^2^, and 0.2 W/cm^2^. From the violin plots, two populations of neurons appear – one with high baseline activity and one with low baseline activity. At power densities of 2.0 W/cm^2^, 1.0 W/cm^2^, and 0.2 W/cm^2^, the more active population disappears or shrinks. Almost all the cells experience a decrease in activity at these power densities, while only around half show a decrease at 0.02 W/cm^2^, which may be attributed to spontaneous fluctuations in activity. Furthermore, the cells with the highest baseline activity experience the greatest decreases in activity. No inhibition was observed at off-resonance frequencies (**Supplementary fig. 2**). The changes in strength of inhibition and radius of the affected area demonstrate a dependence of microwave inhibition on power density.

**Figure 3:**
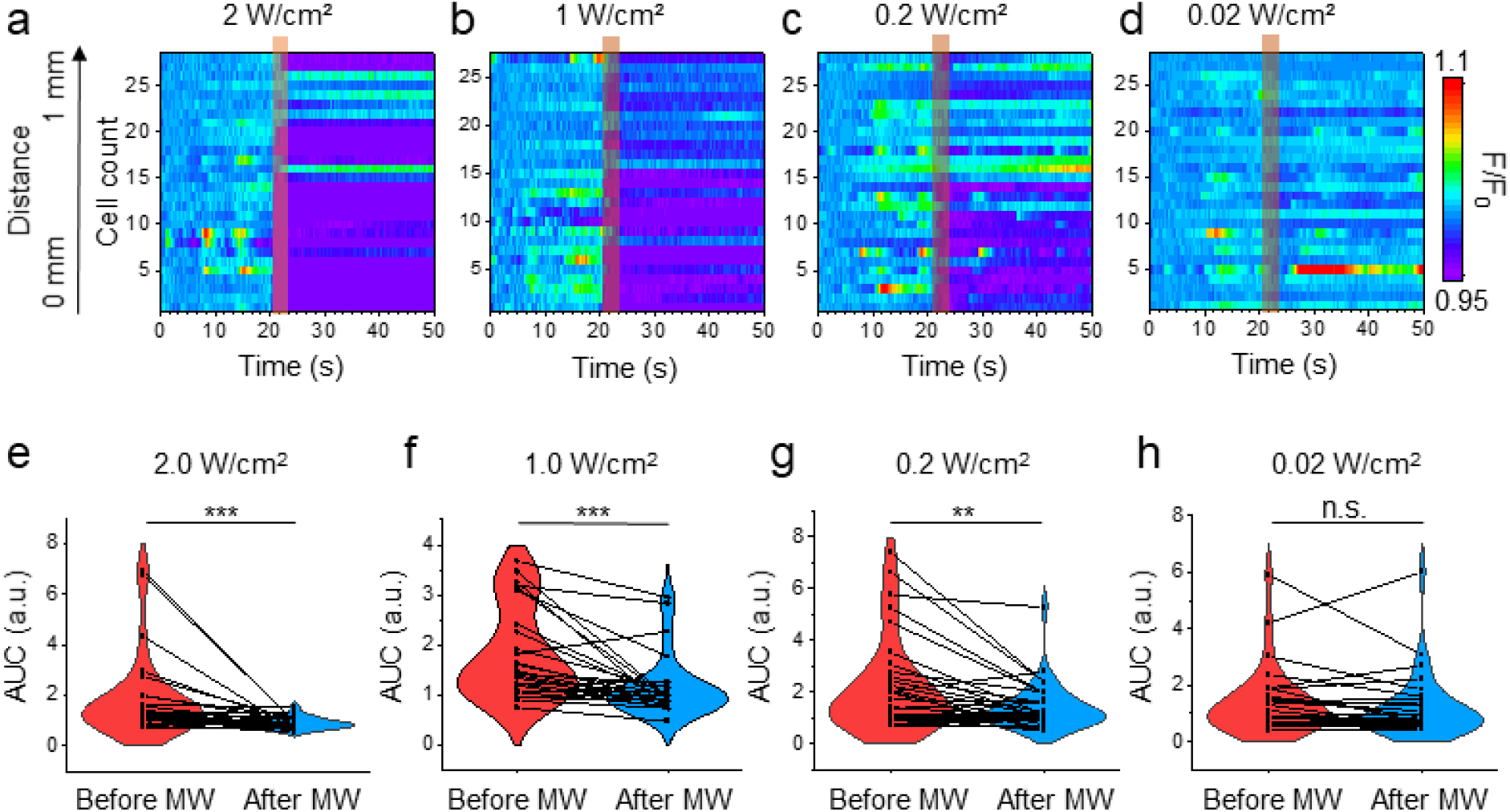
The SRR enhances the efficiency and spatial resolution of microwave inhibition. (a, b, c, d) GCaMP6f fluorescence heatmaps for neurons within 1 mm from the SRR gap under 2.0 GHz microwave for 3 s at varying power densities; cells are arranged by distance from SRR gap (e, f, g, h) AUC of F/F_0_ traces for cells in (a, b, c, d), respectively, before and after microwave irradiation (n=28, n=26, n=34, n=30); Shaded boxes indicate microwave on periods; statistical significance calculated using paired sample Wilcoxon signed rank test where ***p<0.001, **p<0.01, *p<0.05.

To observe how the SRR enhances inhibition efficiency of the microwave, a direct comparison can be made between neurons treated at 0.2 W/cm^2^ with (**Figure 3c**) and without (**Figure 2d**) the SRR. With the SRR, neuronal activity was reduced by 36.3% (**Figure 3g**). Without the SRR, microwave treatment did not induce any significant change in neuronal activity (**Figure 2e**). Furthermore, the fluorescence heatmap of neurons within 1 mm from the SRR gap at 0.2 W/cm^2^ shows that, at this power, the SRR achieves submillimeter spatial precision. Together, these findings demonstrate that the SRR can inhibit neuronal activity with an improved efficiency and spatial precision over direct microwave, enabling much lower microwave dosage to be implemented.

### Biocompatible titanium SRR inhibits neurons with sub-millimeter spatial precision

Although the copper SRR has shown neuronal inhibition with enhanced efficiency and spatial precision, the poor compatibility of copper with tissue hinders its capability in biomedical application. Titanium alloy, on the other hand, has shown excellent biocompatibility with tissue and has seen wide application for tissue implants in the clinics, such as in artificial joints and pacemakers [23]. To test if titanium alloy is a better candidate for neuromodulation, a titanium SRR (TiSRR) with an outer diameter of 3.28 mm, gap of 0.239 mm, height of 0.2 mm and width of 0.85 mm was fabricated (**Figure 4a**). The TiSRR was fabricated from a tapered titanium alloy tube sectioned with electron discharge machining. To verify that the TiSRR has a similar resonance effect as the copper SRR, finite element modeling was performed (**Figure 4b**). The copper and titanium SRRs were modeled with the same geometry as the fabricated TiSRR to directly compare the materials. Both SRRs had peak resonance at 1.9 GHz. As expected, the copper SRR generated an 11.8% greater electric field at the resonant frequency. These results confirm that the TiSRR generates a localized electric field similarly to the copper SRR without a great loss in efficiency.

**Figure 4:**
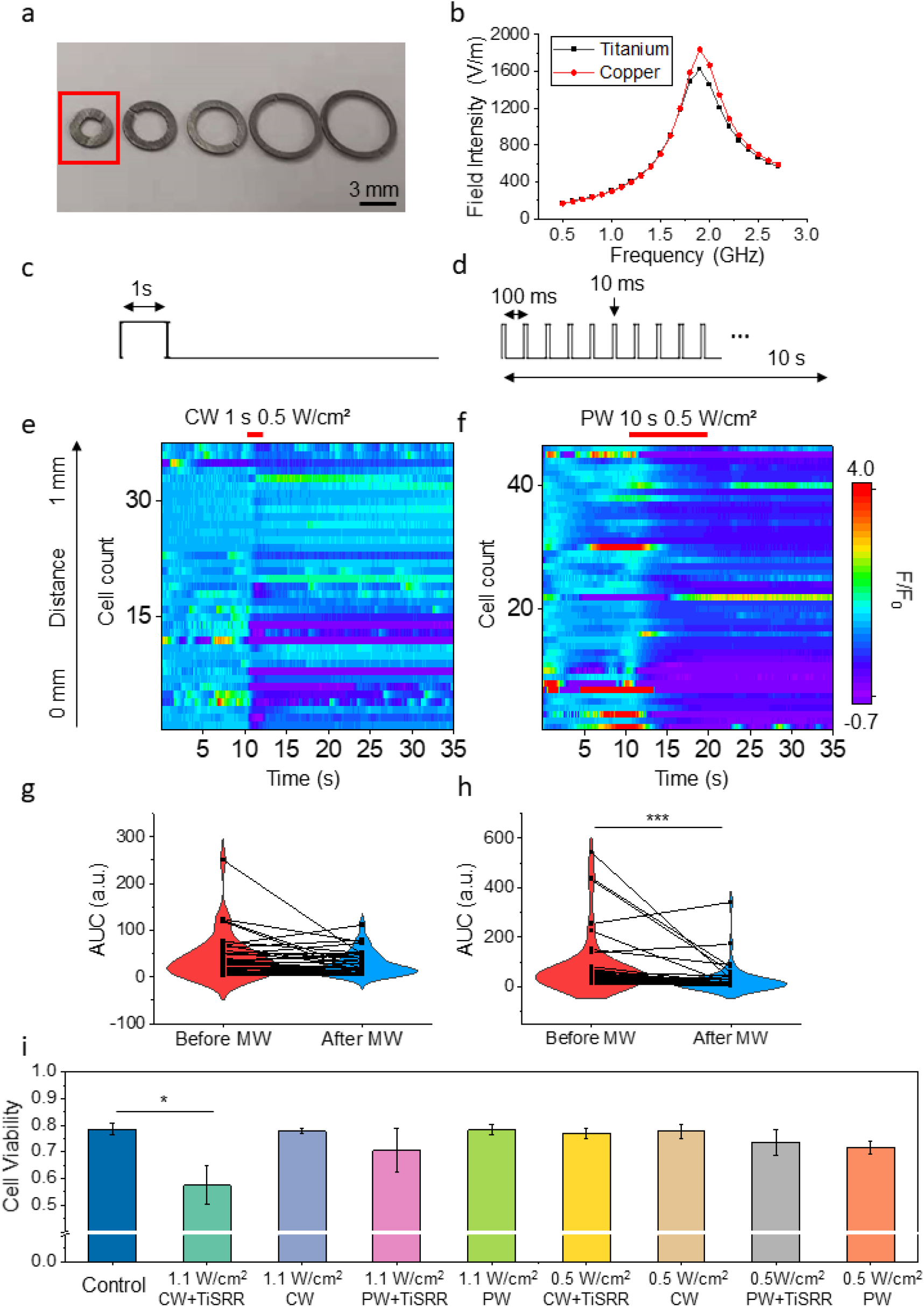
Pulsed microwave improved neuronal inhibition with TiSRR and biocompatibility. (a) TiSRRs were fabricated by sectioning a tapered titanium alloy tube to create SRRs of varying diameters; the TiSRR with 3.28 mm diameter (red box) was chosen; (b) simulation of field intensity at the gap for copper SRR and TiSRR over given frequencies; (c) schematic of continuous microwave profile; (d) schematic of pulsed microwave profile; (e) heatmap of normalized GCaMP6f fluorescence intensity for neurons near the TiSRR gap treated with 1 s continuous microwave at 0.5 W/cm^2^ and 2.1 GHz; (d) same as (e) with pulsed microwave for 10 s; (g, h) AUC of F/F_0_ traces for cells in (e, f), respectively, before and after microwave irradiation (n=37, n=46); statistical significance calculated using paired sample Wilcoxon signed rank test where ***p<0.001, **p<0.01, *p<0.05; (i) cell viability for primary neuron cultures after 10 min of continuous (CW) or pulsed (PW) microwave treatment with or without the TiSRR (n=3, error bars represent standard error of the mean); statistical significance calculated using two sample t test where ***p<0.001, **p<0.01, *p<0.05.

Next, the efficiency of neuronal inhibition with the TiSRR was tested by the same setup as described in **Supplementary fig. 1b**. The resonant frequency of the TiSRR was empirically found to be 2.1 GHz. The discrepancy may be because pure titanium was used in the simulation while the TiSRR was fabricated from titanium alloy. Microwave irradiation of primary cortical neurons with 0.5 W/cm^2^ microwave at 2.1 GHz for 1 s in the presence of the TiSRR demonstrated that the TiSRR achieves neuronal inhibition *in vitro* (**Figure 4e**, **Supplementary fig. 3**). Voltage fluorescence imaging of primary cortical neurons transfected with Archon confirmed that the inhibition effects observed were not artifacts of thermal interference with the GCaMP6f fluorescence (**Supplementary fig. 4**). These results confirm the ability of the TiSRR to inhibit neuronal activity.

For clinical applications, it may be preferable to prolong the microwave inhibition without increasing the thermal accumulation or microwave dosage. To this end, we modulated the microwave to generate a pulse train having a 10% duty cycle over 10 s, i.e., 10 ms pulses with a 10 Hz repetition rate (**Figure 4d**). The overall microwave dosage is equivalent to the 1 s continuous microwave (**Figure 4c**). The continuous and pulsed microwave performance were compared by irradiating primary cortical neurons with 0.5 W/cm^2^ microwave at 2.1 GHz in the presence of the TiSRR (**Figure 4e, f**). While the continuous wave only reduced neuronal activity by 32.9% (**Figure 4g**), the pulsed microwave reduced neuronal activity by 67.6% (**Figure 4h**). The improved performance may be due to a decrease in thermal accumulation, reducing the effects of thermal activation under pulsed microwave. Cell viability was measured after microwave treatment with 1.1 W/cm^2^ and 0.5 W/cm^2^, with or without the TiSRR, and under continuous or pulsed microwave (**Figure 4i**). Treatment, i.e., 1 s continuous or 10 s pulsed microwave, was repeated every 30 s for 10 min. Control neurons were placed at room temperature for 10 min. Only 1.1 W/cm^2^ continuous microwave with the TiSRR had a viability significantly lower than the control. In the case of 1.1 W/cm^2^ with the TiSRR, the pulsed microwave had greater cell viability than the continuous microwave. Taken together, these results indicate that the TiSRR provides a platform for neuronal inhibition with comparable performance to the copper SRR and greater biocompatibility. Furthermore, pulse modulation is a viable method for prolonging microwave treatment without inducing thermal toxicity.

### TiSRR mediates transcranial inhibition of neurons

Our eventual goal is to achieve wireless neuronal inhibition for the treatment of disorders like epilepsy. For the device to be wireless, microwave must be delivered from outside the skull to the implanted SRR. The millimeter-scale wavelength of microwave allows for deep penetration into biological tissue, including bone. Microwave has been demonstrated to penetrate >50 mm into the human skull while maintaining over 50% of its energy [14], making wireless transcranial microwave inhibition feasible.

To evaluate the transcranial inhibition capabilities of the microwave TiSRR, a macaque monkey skull with ~3 mm thickness was placed over the neuron culture inside the microwave waveguide (**Figure 5a**). The TiSRR was placed over primary cortical neurons and irradiated with 0.5 W/cm^2^ pulsed microwave at 2.1 GHz for 10 s (**Figure 5b, c**). With the skull, the TiSRR achieved 52.2% reduction in neuronal activity, while without the skull it achieved 48.7% reduction during microwave treatment (**Figure 5d, e**). As expected, the TiSRR performance was not significantly hindered by the presence of a skull. This result demonstrates the capability of the TiSRR to perform transcranial neuronal inhibition for wireless application of the device.

**Figure 5:**
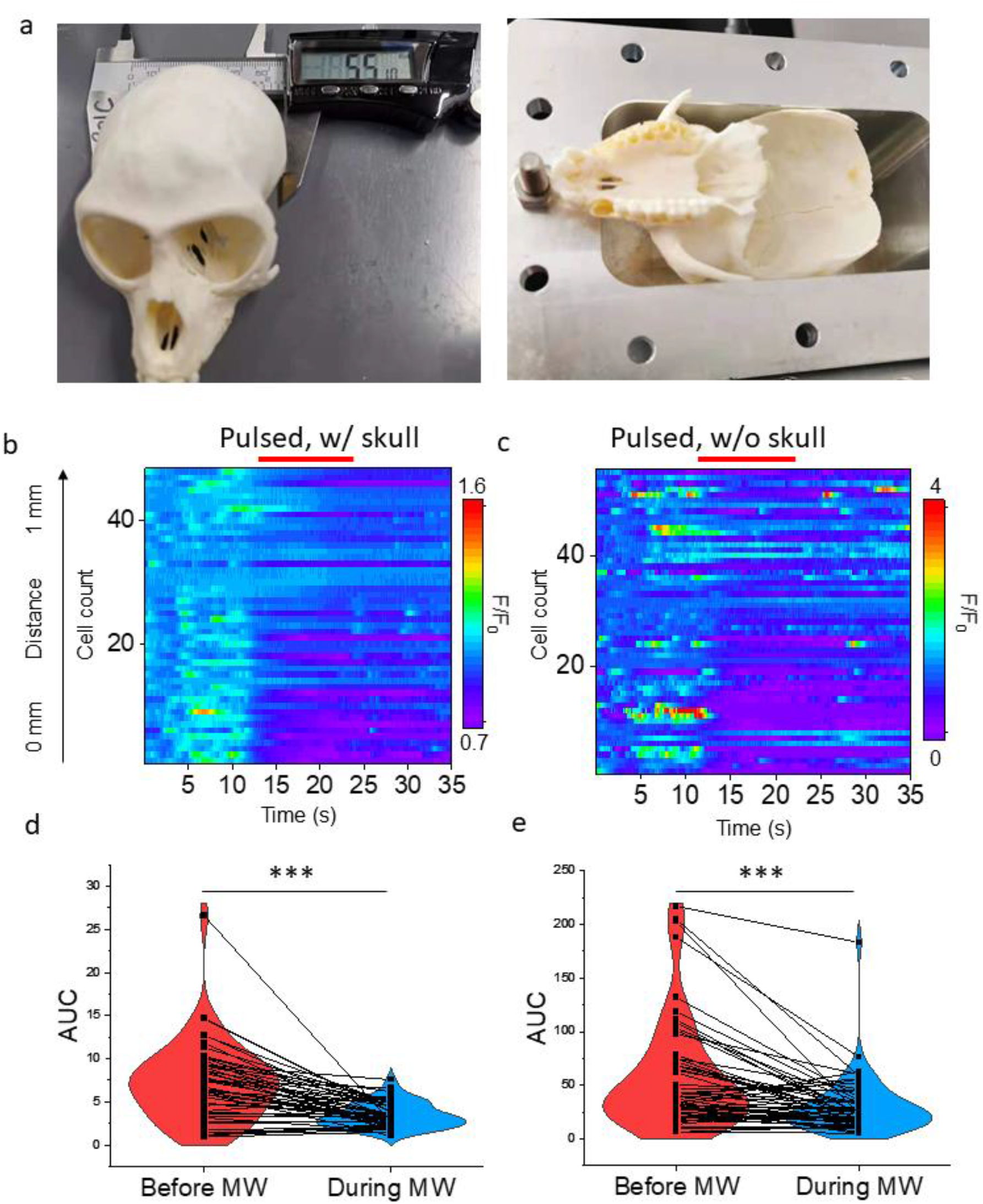
The TiSRR performs transcranial inhibition. (a) A macaque monkey skull with thickness ~3 mm inside the microwave waveguide used for transcranial inhibition; (b, c) GCaMP6f normalized fluorescence heatmap for neurons near the TiSRR gap under 0.5 W/cm2 pulsed microwave irradiation at 2.1 GHz for 10 s with (b) and without (c) the monkey skull; (d, e) AUC of normalized fluorescence intensity for neurons in (b, c), respectively (n=48, n=55); statistical significance calculated using paired sample Wilcoxon signed rank test where ***p<0.001, **p<0.01, *p<0.05.

### TiSRR inhibits stimulated neurons

Epileptic seizures are characterized by excessive neuronal excitability. The kainic acid (KA) mouse model of epilepsy is commonly used to study the disorder. KA is an analog of glutamate that acts as an agonist to kainite receptors. In small doses, KA increases excitability of a cell population. When injected intracerebrally or systemically, KA evokes acute as well as chronic seizures [24]. To demonstrate the ability of the TiSRR to inhibit KA-induced activity, 20 mM KA in DMSO was added to primary cortical neurons. Treatment with KA noticeably increased the fluorescence intensity (**Figure 6a, b**). The TiSRR was placed over the neurons and irradiated with 0.5 W/cm^2^ pulsed microwave at 2.1 GHz for 10 s (**Figure 6c, d**). Neuronal inhibition was evident, with a 16.3% reduction in normalized fluorescence intensity (**Figure 6f**). For most cells, inhibition only occurred during microwave irradiation, but for some, the effect was longer lasting (**Figure 6e**). The immediate return in activity supports a non-toxic mechanism for microwave inhibition of neurons. The inhibitory effect was confined to ~200 μm from the SRR gap (**Figure 6d**), further demonstrating the submillimeter spatial precision of the TiSRR. These results indicate that the TiSRR can inhibit a stimulated increase in neuronal activity with high spatial precision.

**Figure 6:**
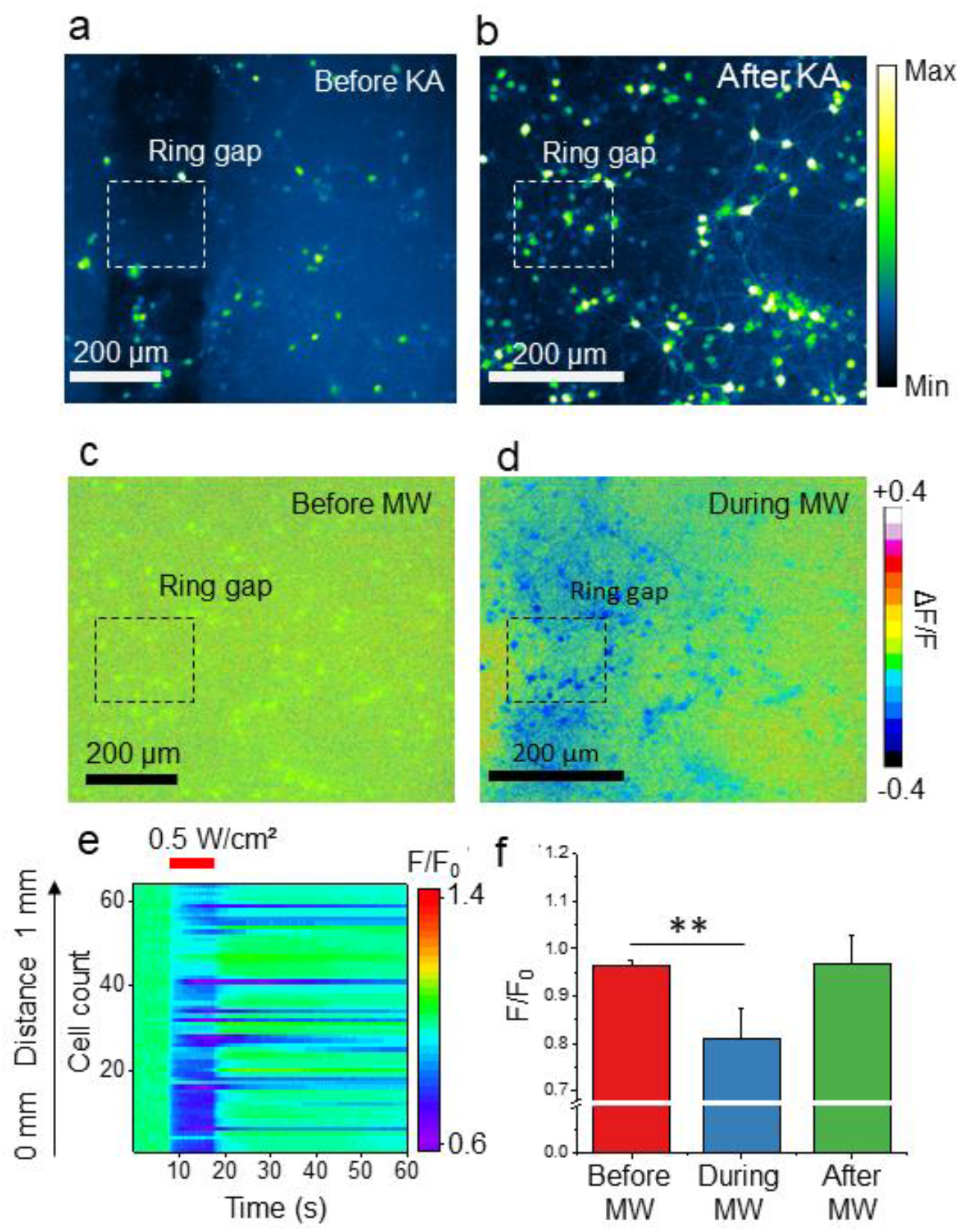
The TiSRR inhibits kainic acid-induced activity *in vitro*. (a, b) GCaMP6f fluorescence images of neurons before (a) and during (b) treatment with 20 mM kainic acid; (c, d) ΔF/F normalized images of (a, b), respectively; (e) GCaMP6f normalized fluorescence heatmap of kainic acid-induced activity suppressed by the TiSRR with 0.5 W/cm^2^ pulsed MW at 2.1 GHz for 10 s; (f) bar graphs of normalized fluorescence intensity for cells in (e) (n=64, error bars represent standard deviation); statistical significance calculated using two sample t test where ***p<0.001, **p<0.01, *p<0.05.

### TiSRR suppresses seizures in mouse model of epilepsy without tissue damage

As a further proof-of-concept for epilepsy treatment with the microwave TiSRR, a mouse model of epilepsy was used. Picrotoxin is a GABA antagonist and thus a convulsant. Male C57BL/6J mice aged 14-16 weeks were injected with picrotoxin to induce seizure. The cortex was chosen as the seizure target to avoid implantation of the TiSRR, which is too large for the mouse brain. A localized seizure was induced in the cortex by injecting 10 nL of 20 mM picrotoxin in DMSO. EEG recordings were taken at the injection site. The TiSRR was placed on the surface of the cortex with the gap directly on the injection site. Because the resonant frequency of the TiSRR is dependent upon the medium it is in, the TiSRR was covered in ultrasound gel, which has properties similar to water. Microwave treatment consisting of 10 s of pulsed microwave at 2.1 GHz and 0.5 W/cm^2^ was applied (**Figure 7a**). After microwave treatment, the average spike amplitude significantly reduced (**Figure 7b**). These results demonstrate that the TiSRR has the ability to reduce neuronal activity *in vivo*. With optimized microwave treatment parameters, the TiSRR may be able to further reduce the magnitude, and thus the impact, of a seizure.

**Figure 7:**
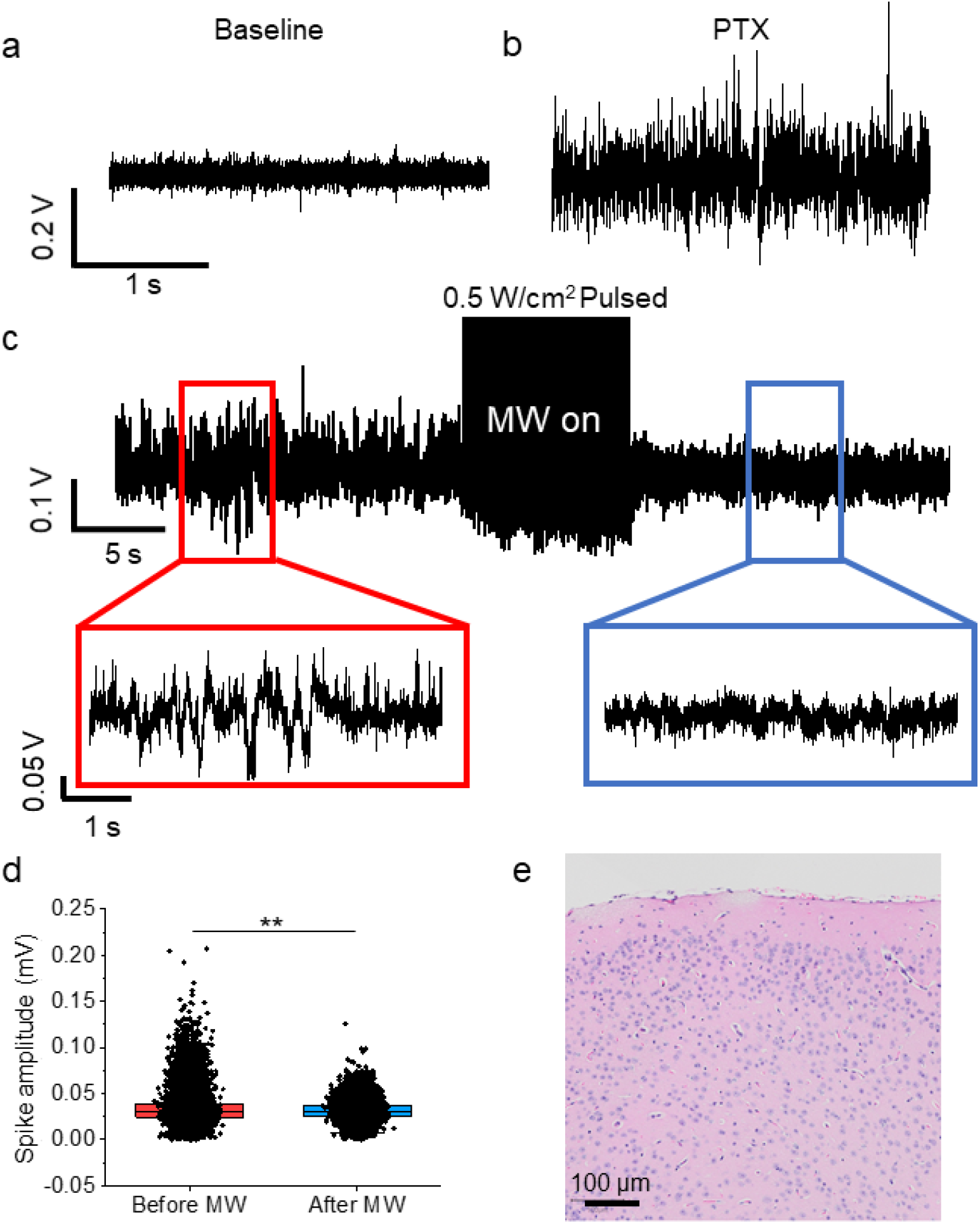
TiSRR reduces picrotoxin-induced activity *in vivo*. (a) Baseline EEG recording of mouse cortical neurons; (b) EEG recording of mouse cortical neurons after intracerebral injection of 10 mM picrotoxin; (c) TiSRR was placed over the injection site and irradiated with 0.5 W/cm^2^ pulsed microwave at 2.1 GHz for 10 s; (d) spike amplitude before and after microwave treatment with TiSRR; (e) brain tissue treated with 10 s pulsed microwave at 0.5 W/cm^2^ and 2.1 GHz for 6 min and stained with H&E; statistical significance calculated using two sample t test where ***p<0.001, **p<0.01, *p<0.05.

Next, we evaluated the safety of our microwave treatment for *in vivo* applications. A mouse brain was exposed to 10 s of pulsed microwave at 2.1 GHz and 0.5 W/cm^2^ once every minute for 6 minutes. This treatment was chosen to demonstrate a higher overall dosage than the inhibition conditions while staying within the safe exposure limits. The cortical tissue was paraffin embedded, sectioned, and H&E stained. The histology appeared normal, showing no signs of hemorrhage or cell death (**Figure 7c**). These results indicate that the maximum microwave dosage that would be used *in vivo* does not induce damage in brain tissue.

## Discussion

This work presents wireless neuromodulation at sub-millimeter precision enabled by a tailor-designed microwave split-ring resonator. Microwave has not previously been used to modulate neurons *in vivo* because at high power it can cause thermal damage. As demonstrated here, by implanting an SRR in the deep brain, microwave inhibition efficiency is much improved and dosages below the safe exposure limit can be used.

Wirelessly powered neural implants have received great attention in recent years. These implants possess clear advantages over tethered devices in that they reduce tissue damage during surgical procedures and, subsequently, diminish infection in daily use. However, a primary challenge for wireless neural stimulators is to create efficient miniature devices that operate at deep tissue. For efficient wireless power transfer, antennas need to have sizes comparable to the electromagnetic wavelength. Currently, the majority of miniaturized wireless neural modulators work in the MHz range and require a surface-level receiver to couple with the waves to reach the deep brain, increasing the invasiveness and size of the implant [25, 26]. For fully internalized devices, power delivery becomes difficult due to their small size, thus limiting the depth of the implants [27]. More recently, ultrasound-powered neural modulators have enabled effective power transfer at several centimeters deep into the tissue [28]. Such devices, however, are difficult to operate in free moving animals due to the impedance mismatch between air and soft tissue, thus requiring direct contact and application of ultrasound gel. Compared to these devices, our SRR offers several unique advantages (**Table 1**). First, the SRR creates a microwave field with ultrahigh spatial precision on the order of 100 μm, which is one-hundredth the wavelength of microwave. This precision enables region-specific brain modulation or selective inhibition of a single nerve. Second, the implantable, miniaturized SRR has a volume of 1.8 mm^3^, which makes it the smallest implant for wireless neuromodulation. This small size greatly reduces invasiveness and minimizes the wound healing response. Third, the SRR allows wireless neural inhibition at centimeter-scale depths. This capability enables deep-tissue modulation for the treatment of disorders involving excessive excitability, such as neuropathic pain.

**Table 1.** Comparison of microwave SRR with existing neuromodulation implants.

| Device name | Power type | Implant size ($\text{mm}^3$) | Free-moving | Modulation type | Penetration depth |
| --- | --- | --- | --- | --- | --- |
| Micro LED [29, 26] | Radio frequency | 25-50 | Y | excitation | Several cm |
| Stim Dust [27] | Piezoelectric | 10 | N | excitation | Several cm |
| Magneto-electric stimulator [28] | Magnetic field | 175-500 | Y | excitation | Several cm |
| Microwave SRR (this work) | Microwave (1.0-2.5 GHz) | 1.8 | Y | inhibition | Several cm |

A major advantage of the SRR is that it allows the use of microwave dosages within the safety limits established by IEEE [20]. The threshold for safe RF exposure is 10 W/kg averaged over 6 minutes, which corresponds to an average dosage of 3600 J/kg. Each treatment, consisting of 10 s pulsed microwave at 0.5 W/cm^2^, corresponds to 500 J/kg in vitro (17.5 mm radius, 5 mm depth). This means we can administer up to 7 sessions of treatment within 6 minutes according to IEEE standards. Our dosage is also below those used in previous literature [25]. Furthermore, the major mechanism behind microwave toxicity is thermal damage to the blood brain barrier (BBB) [20, 30, 31]. Ikeda et al. found that the dog brain could withstand temperatures up to 42°C for 45 min before irreversible damage to the BBB occurred [31]. Studies in other species – including rats, monkeys, rabbits, and pigs – revealed that most brains could withstand at least 1 min at 43°C without damage, with pig brains lasting over 150 hours [32, 33]. When placed in bulk PBS and irradiated with 10 s pulsed microwave at 0.5 W/cm^2^, the SRR gap reached a peak of 24°C (22°C baseline). Therefore, our device operates within the safety parameters for microwave exposure to the brain.

While microwave inhibition of neurons has previously been established, its mechanism remains unclear. It has been proposed that millimeter waves (MMW) modulate neurons via a thermal mechanism [34], but our work and others suggest the effect may be more complex [35, 36, 15]. While neurons are modulated by changes in temperature, thermal increase is known to stimulate neurons rather than inhibit them by increasing the energy of molecules involved in channel gating [34]. It has been observed that thermal increase alone cannot reproduce all of the effects of MMW in excitable cells, such as induction of nitric oxide production [35] and suppression of firing rate [36, 15]. In this work, we found the pulsed microwave, which induces less thermal increase than equivalent dosage of continuous microwave, to be more effective for inhibition. It is further worth noting that at power densities of ~3 W/cm^2^, we observed thermal stimulation of neurons via the SRR (not shown), suggesting that inhibition occurs by a separate mechanism which is overcome by thermal effects at certain power densities.

Our evidence suggests that a nonthermal mechanism contributes to microwave inhibition of neurons. However, the detailed biophysical mechanism is still unclear. There are several ways in which inhibition might occur. The rotational spectrum of water lies in the 5 to 125 cm^-1^ range [37], which overlaps with the frequency range of microwaves. This likely implicates altered behaviors of water molecules at the membrane-water interface [38, 39] and/or crevices in ion channels [40, 41] upon microwave irradiation. In addition, perturbation in interfacial water properties is also expected to affect the solvation of ions, also leading to changes in ion channel activities [42, 43]. These effects can potentially hyperpolarize the cell and increase the cell’s activation threshold. Future work will investigate this possible mechanism of microwave inhibition. An understanding of the mechanisms behind microwave inhibition of neurons will enable more efficient and safer implementation of the device.

The microwave SRR is a novel platform for wireless, battery-free neuromodulation in the deep brain with high spatial precision. The device operates within safety limits and occupies a volume < 2 mm^3^. Improvements to the device might include making the implant a rod so that it may be injected into the brain. In future work, the SRR may be applied to other conditions, such as chronic pain or Parkinson’s Disease. Multiple SRRs with varying diameter may be implanted to modulate multiple brain regions in sequence. The thermal stimulation capabilities of the SRR at power densities around 3 W/cm^2^ will also be investigated for use on its own or in conjunction with inhibition.

## Methods

### Numerical simulation of the resonant frequency of SRR

Simulations were performed in COMSOL Multiphysics 5.3a. All SRRs were modeled in bulk water medium with electrical conductivity 5.5×10^-6^ S/m and a relative permittivity of 70. Material parameters were taken from the copper model and the solid beta-phase titanium model in the COMSOL materials library. The microwave originated from a 50 cm^2^ port with a plane wave input that has *E* polarized in the *y*-direction. *H* was polarized perpendicular to the SRR plane in the *z*-direction. Scattering conditions were used at the boundaries of the simulated area.

### Copper SRR fabrication

The copper SRR was laser cut from a copper sheet by Kuso-Relock USA LLC.

### Titanium SRR fabrication

The TiSRR was fabricated from a titanium alloy tube with outer diameter tapering from 2 mm to 4 mm. Electrical discharge machining (EDM) wire cutting with a 100-μm diameter wire was used to create a slit of 200 μm down the length of the tube. Then, multiple parallel cuts were made every 200 μm perpendicular to the slit to produce SRRs of varying diameters.

### Cell culture

Primary cortical neurons were harvested from Sprague-Dawley rats at embryonic day 18 (E18). Cortices were dissected from rats of either sex and digested with papain (0.5 mg/mL in Earle’s balanced salt solution) (Thermofisher Scientific). Neurons were plated onto poly-D-lysine coated glass bottom culture dishes in Dulbecco’s Modified Eagle Medium (Thermofisher Scientific) with 10% fetal bovine serum (Thermofisher Scientific). After 24 hours, medium was replaced with feeding medium consisting of Neurobasal medium supplemented with 2% B-27 (Thermofisher Scientific), 1% N2, and 1% GlutaMAX™ (Thermofisher Scientific). 0.1% 5-fluorodeoxyuridine (FdU) was also added to remove glial cells. At this time point, neurons were incubated with 0.1% pAAV.Syn.Flex.GCaMP6f.WPRE.SV40 (Addgene) for calcium imaging or pAAV-Syn-Archon1-KGC-GFP-ER2 (Addgene) for voltage imaging. Fresh feeding medium was added to the culture every 3-4 days.

### Thermal imaging

The SRR was placed in a plastic dish and immersed in PBS. The microwave waveguide was oriented with *H* field perpendicular to the ring plane. Microwave was delivered for 1 s at the resonance frequency and 2 W/cm^2^. Imaging was performed using a thermal camera (A325sc, FLIR). Video was captured at a frame rate of 30 Hz for 10 s.

### Calcium imaging

Calcium imaging was performed on a lab-built microscope based on an Olympus IX71 microscope frame with a 20x air objective (UPLSAPO20X, 0.75 NA, Olympus). The sample was illuminated by a 470 nm LED (M470L2, Thorlabs), with an emission filter (FBH520-40, Thorlabs), an excitation filter (MF469-35, Thorlabs), and a dichroic mirror (DMLP505R, Thorlabs). A scientific CMOS camera (Zyla 5.5, Andor) was used to collect images at 20 frames per second.

### Voltage imaging

Voltage imaging was performed on a lab-built microscope based on an Olympus IX71 microscope with a 20x air objective (UPLSAPO20X, 0.75 NA, Olympus). The sample was illuminated by a 638 nm excitation laser (0638-06-01-0180-100, Cobolt) with a dichroic beam splitter (Di01-R405/488/532/635, Semrock), a 675 nm emission filter (Chroma), and a 695 nm emission filter (HPM695-50, Newport). A digital CMOS camera (C13440, Hamamatsu) was used to collect images at 400 frames per second.

### Animal surgery

All experimental procedures have complied with all relevant guidelines and ethical regulations for animal testing and research established and approved by the Institutional Animal Care and Use Facility of Boston University under protocol PROTO201800534. C57BL/6J mice aged 14-16 weeks were anaesthetized using 5% isoflurane in oxygen then maintained with 1.5-2% isoflurane via nose cone throughout the procedure and experiment. Tail pinch was used to monitor anaesthetization throughout, and body temperature was maintained with a heat pad. The hair and skin on the dorsal surface were removed. A craniotomy was performed using a dental drill to remove a ~3 mm diameter patch of skull over the right hemisphere. Saline was applied to immerse the brain. In relevant experiments, the TiSRR was placed on the cortical surface over the injection site and covered in ultrasound gel. After seizure induction and microwave treatment, mice were perfused with saline and 10% formalin. The brain was removed, paraffin embedded, sectioned, and H&E stained at the Boston University Collaborative Research Laboratory. Imaging was performed in the Boston University BME Micro/Nano Imaging Facility.

### Seizure induction and electrocorticogram recording

A modified version of the procedure described by Melo-Carillo et al. was used [48]. Seizure was chemically induced by injecting 10 nL of 20 mM picrotoxin in DMSO into the cortex at AP-2, ML+2, DV+0.5, where bregma was calibrated to be coordinate (0,0). Picrotoxin was injected using a motorized stereotaxic system (Stoelting) at a rate of 5 nL/min. The needle was kept in place for 2 min after injection. A tungsten microelectrode (0.5 to 1 MΩ, Microprobes) was inserted for LFP recording at the injection site. Extracellular recordings were acquired using a Multiclamp 700B amplifier (Molecular Devices) filtered at 0.1 to 100 Hz, digitized with an Axon DigiData 1550 digitizer (Molecular Devices), and denoised with a D400 Multi-channel 60Hz Mains Noise Eliminator. For quantification of response to treatment, a 10 s period of seizure activity before microwave irradiation was used to obtain a baseline mean and SD. Spike amplitude was used to measure neuronal response.

### MW treatment

Microwave was generated using a microwave signal generator (9 kHz to 3 GHz, SMB100A, Rohde & Schwarz) connected to a solid-state power amplifier (ZHL-100W-242+, Mini Circuits) to amplify the microwave to 100 W peak power. Microwave was delivered from a 50 cm^2^ waveguide (WR430, Pasternack) oriented with *H* field perpendicular to the SRR at the resonance frequency of the SRR. The waveguide was ~2 cm from the SRR. Pulse modulation was achieved using a function generator (33220A, Agilent). *In vivo*, one round of treatment consisted of 10 s of 0.5 W/cm^2^ microwave with pulse width 10 ms and repetition rate 10 Hz.

### Data analysis

Calcium images were analyzed using ImageJ. The somata of neurons were selected for fluorescence measurement. Calcium traces, temperature traces, and electrophysiological traces were analyzed using Origin 2018. Area under the curve was calculated by finding the baseline fluctuations with the msbackadj function in the Matlab Bioinformatics toolbox and subtracting the baseline from the fluorescence trace (**Supplementary fig. 6)**. Statistical significance was calculated using paired sample Wilcoxon signed rank test or two sample student’s t test.

## Acknowledgements

This work is supported by NIH R01 NS109794 and R21 EY034275 to JXC and CY; NSF NRT: National Science Foundation Research Traineeship Program (NRT): Understanding the Brain (UtB): Neurophotonics DGE-1633516 to CM; ARO W911NF2110132 to CY; and the Department of Defense (DoD) through the National Defense Science & Engineering Graduate (NDSEG) Fellowship Program to CM. Research reported in this publication was supported by the Boston University Micro and Nano Imaging Facility and the Office of the Director, National Institutes of Health of the National Institutes of Health under award Number S10OD024993. The content is solely the responsibility of the authors and does not necessarily represent the official views of the National Institute of Health. The authors thank Dr. Alexander Rotenberg for his guidance on the epilepsy model.

## Author contributions

CM and YJ carried out the experiments and wrote the manuscript. YL and NZ helped in SRR characterization. LL provided guidance in SRR design. CY and JXC provided overall guidance and manuscript revision.

## Data Availability

The authors declare that the main data supporting the findings of this study are available within the article and its Supplementary Information files. Extra data are available from the corresponding author upon request.

## Code Availability

Data processing code related to the work is available upon reasonable request to the corresponding author.

---

**Supplementary figure 1:**
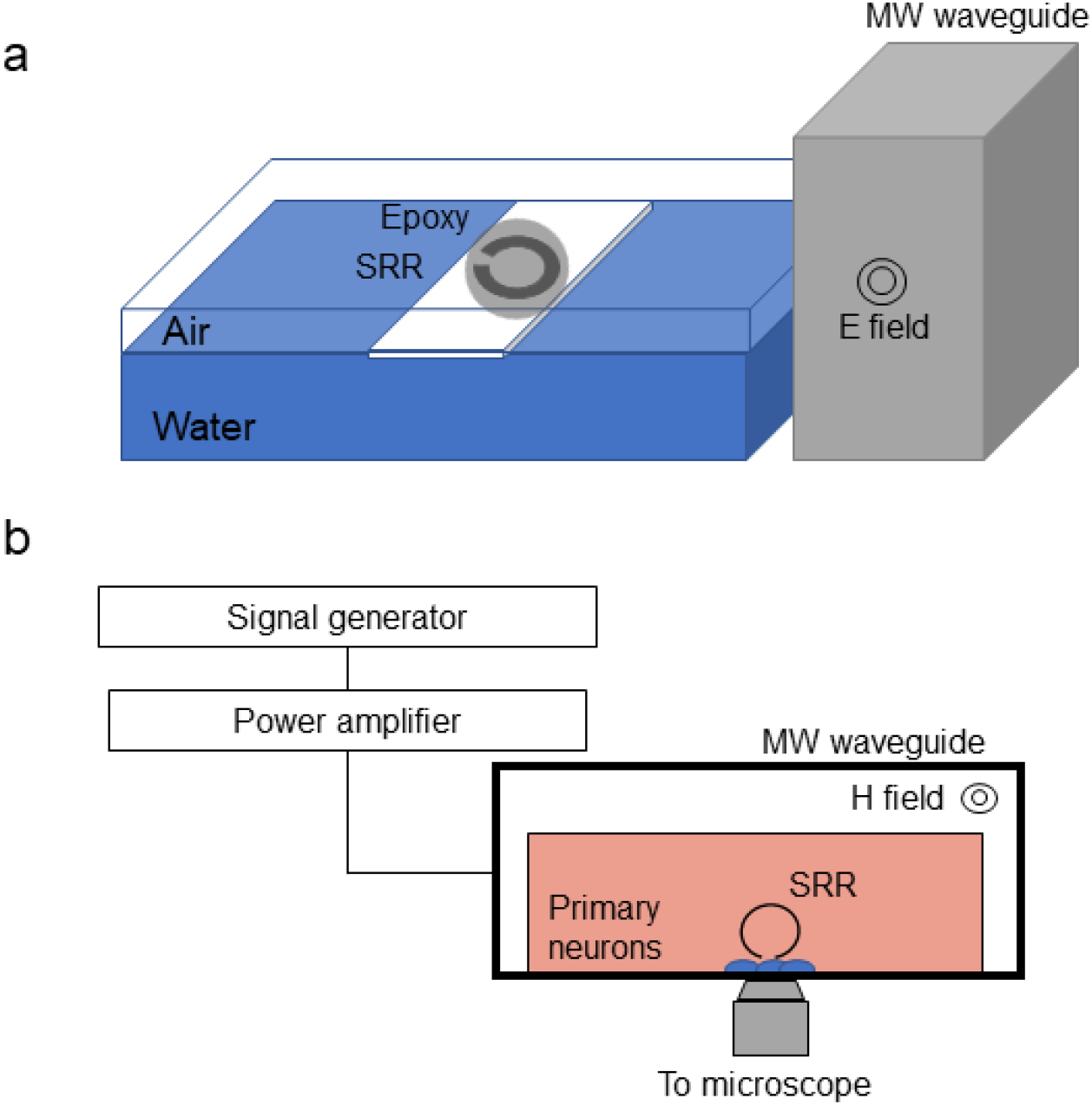
Experimental setup. (a) An SRR with a diameter of 2.54 mm was covered in epoxy, placed at the air-water interface, and irradiated with 2 W/cm^2^ microwave at 2.0 GHz for 1 s; (b) the SRR was submerged in the culture medium above the primary cortical neurons at a distance of ~100 μm from the cells and oriented perpendicular to the culture dish; microwave was delivered with H field perpendicular to the SRR plane.

**Supplementary fig 2.**
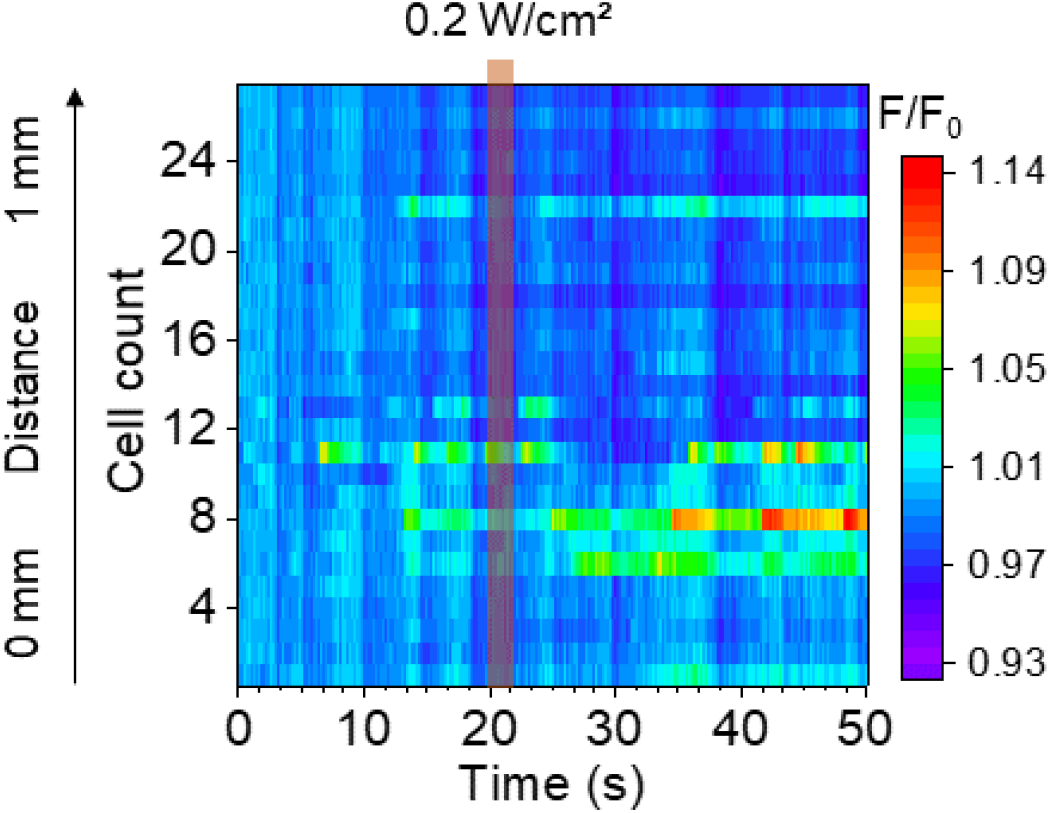
SRR does not induce inhibition at off resonance frequency. GCaMP6f fluorescence heatmap for neurons within 1 mm from the SRR gap under 1.2 GHz microwave for 3 s at 0.2 W/cm^2^.

**Supplementary figure 3:**
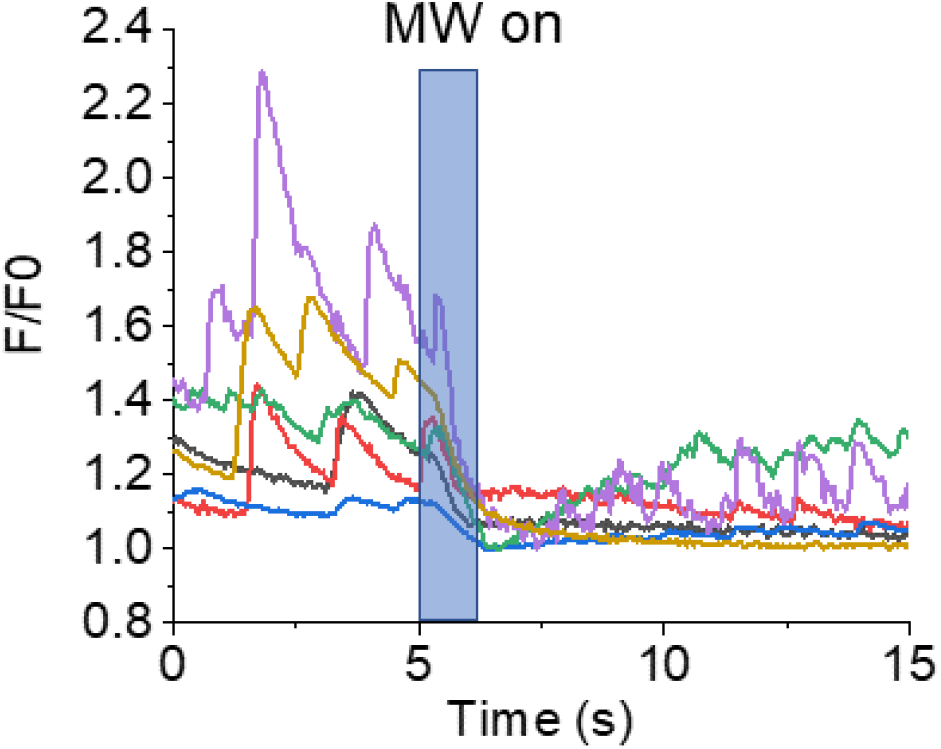
The TiSRR inhibits neuronal activity. GCaMP6f fluorescence traces for neurons near the TiSRR gap irradiated with 0.5 W/cm^2^ microwave at 2.1 GHz for 1 s.

**Supplementary figure 4:**
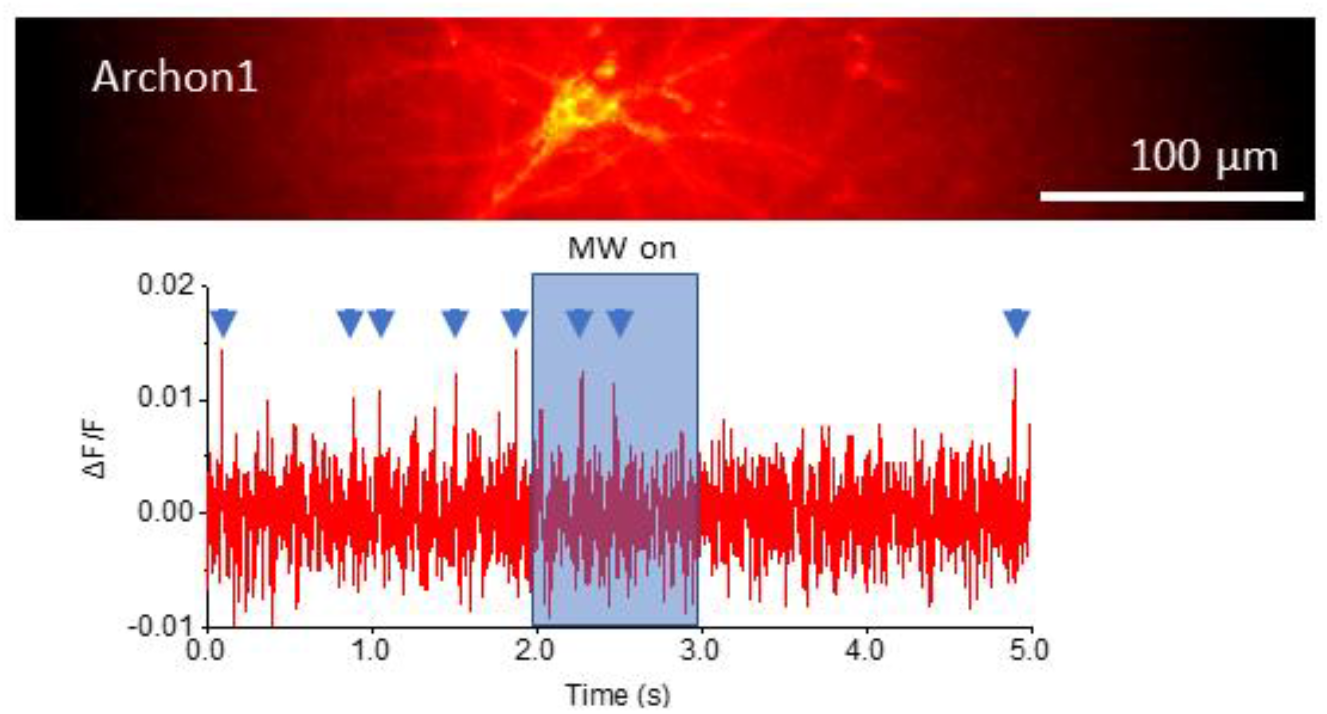
TiSRR inhibition is confirmed by voltage fluorescence imaging. Neurons transfected with Archon1 irradiated with 2 W/cm^2^ microwave at 2.1 GHz for 1 s; blue arrows indicate action potentials.

**Supplementary figure 5:**
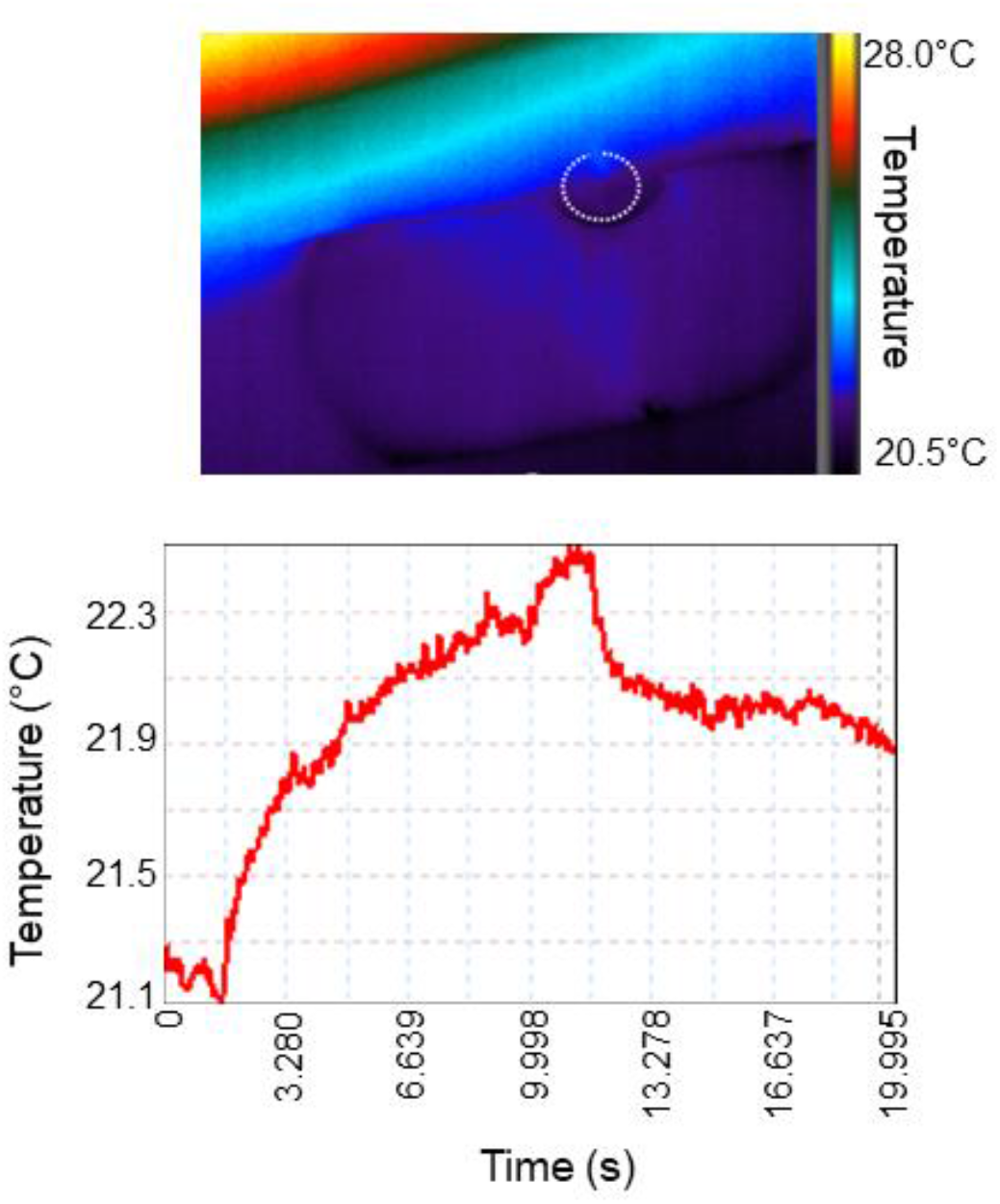
Thermal imaging of SRR under 0.5 W/cm^2^ pulsed microwave.

**Supplementary fig. 6:**
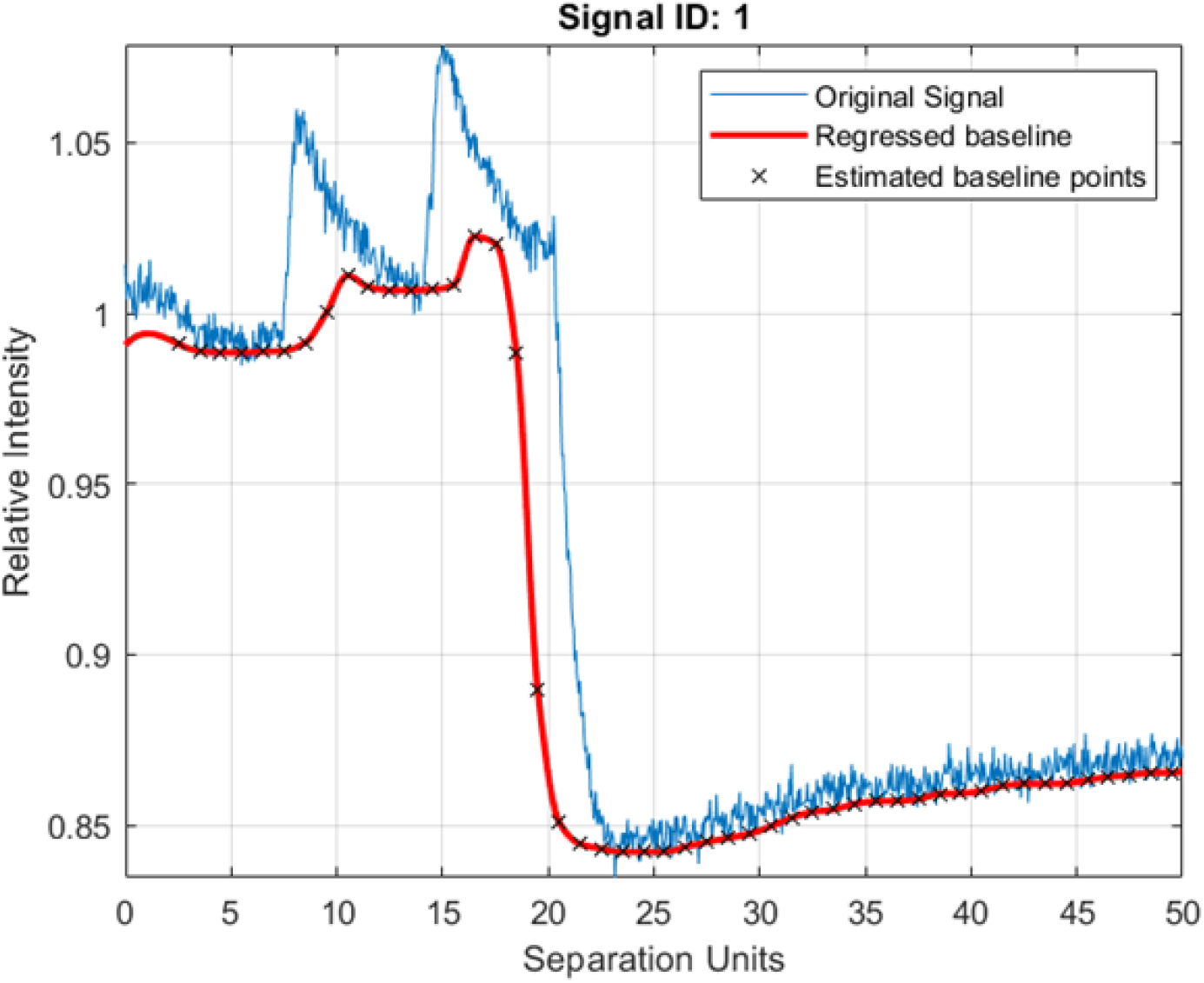
Example of baseline calculation using the msbackadj function from the Matlab Bioinformatics Toolbox.

